# Limited dispersion and quick degradation of environmental DNA in fish ponds inferred by metabarcoding

**DOI:** 10.1101/459321

**Authors:** Jianlong Li, Lori-Jayne Lawson Handley, Lynsey R. Harper, Rein Brys, Hayley V. Watson, Bernd Hänfling

## Abstract

Environmental DNA (eDNA) metabarcoding is a promising tool for rapid, non-invasive biodiversity monitoring. In this study, eDNA metabarcoding is applied to explore the spatial and temporal distribution of eDNA in two ponds following the introduction and removal of two rare fish species. When two rare species were introduced and kept at a fixed location in the ponds, eDNA concentration (i.e., proportional read counts abundance) of the introduced species typically peaked after two days. Thereafter, it gradually declined and stabilised after six days. These findings are supported by the highest community dissimilarity of different sampling positions being observed on the second day after introduction, which then gradually decreased over time. On the sixth day, there was no longer a significant difference in community dissimilarity between sampling days. The introduced species were no longer detected at any sampling positions 48 hrs after removal from the ponds. The eDNA signal and detection probability of the introduced species were strongest near the keepnets, resulting in the highest community variance of different sampling events at this position. Thereafter, the eDNA signal significantly decreased with increasing distance, although the signal increased slightly again at 85 m position away from the keepnets. Collectively, these findings reveal that eDNA distribution in lentic ecosystems is highly localised in space and time, which adds to the growing weight of evidence that eDNA signal provides a good approximation of the presence and distribution of species in ponds. Moreover, eDNA metabarcoding is a powerful tool for detection of rare species alongside more abundant species due to the use of generic PCR primers, and can enable monitoring of spatial and temporal community variance.

## 1 Introduction

Environmental DNA (eDNA) analysis has emerged as a powerful tool in biological conservation for rapid and effective biodiversity assessment. This tool relies on the detection of genetic material that organisms leave behind in their environment (Taberlet et al. 2012; Thomsen & Willerslev 2015). An important application of this method is discovery, surveillance and monitoring of invasive, rare, or threatened species, especially in environments where organisms or communities are difficult to observe, such as aquatic environments (reviewed in Rees et al. 2014; Lawson Handley 2015; Barnes & Turner 2016; Deiner et al. 2017). Several studies have found positive relationships between eDNA concentration and organism density in aquatic ecosystems (e.g., Takahara et al. 2012; Pilliod et al. 2013; Li et al. 2018a). However, in freshwater ecosystems, the detection probability of eDNA is highly dependent on its characteristics, including the origin (physiological sources), state (physical forms), transport (physical movement), and fate (degradation) of eDNA molecules (reviewed in Barnes & Turner 2016). Consequently, the understanding of eDNA characteristics is crucial to improve eDNA sampling designs and ensure the accuracy and reliability of eDNA biodiversity assessments (Goldberg et al. 2018).

Organisms shed DNA into their environment as sloughed tissues (e.g., faeces, urine, moulting, mucus or gametes) and whole cells, which then break down and release DNA (reviewed in Lawson Handley 2015; Thomsen & Willerslev 2015). Studies have demonstrated that eDNA production rates can be highly variable among species in aquatic ecosystems (Goldberg et al. 2011; Thomsen et al. 2012b; Sassoubre et al. 2016), and several factors can influence the amount of genetic material released by organisms into water, including biomass, life stage, breeding, and feeding behaviour (Maruyama et al. 2014; Pilliod et al. 2014; Klymus et al. 2015; Tillotson et al. 2018).

Once released into the environment, eDNA is transported away from organisms and begins to degrade. To better understand the distribution of eDNA in relation to species distribution, investigations have begun to examine how this complex DNA signal is transported horizontally (i.e., downstream) and vertically (i.e., settling) in aquatic environments. In lotic ecosystems, including rivers and streams, eDNA studies on horizontal transport produced variable results, where eDNA is transported metres to kilometres depending on stream discharge (Deiner & Altermatt 2014; Pilliod et al. 2014; Jane et al. 2015; Jerde et al. 2016). In contrast to lotic ecosystems, the natural hydrology of lentic ecosystems, such as lakes and ponds, may be less complex. In still water, eDNA has been shown to accumulate nearby to target organisms, with detection rate and eDNA concentration dropping off dramatically less than a few metres from the target organisms (Takahara et al. 2012; Eichmiller et al. 2014; Dunker et al. 2016). Additionally, eDNA detection may provide a more contemporary picture of species distribution, as transport is less important in lentic ecosystems. This may allow for greater settling of eDNA in sediment at the location where DNA shedding took place. Indeed, eDNA concentration of targeted fish is higher in sediment than in surface water of lentic systems (Eichmiller et al. 2014; Turner et al. 2015). Therefore, sedimentary eDNA can also result in false positive detections and affect inferences made regarding the current presence of a species.

eDNA degradation can also reduce the detectability of species over time. The rate of degradation in water can range from hours to weeks, depending on the ecosystem, target species, and eDNA capture method in question (Dejean et al. 2011; Takahara et al. 2012; Thomsen et al. 2012a; Thomsen et al. 2012b; Goldberg et al. 2013; Balasingham et al. 2017; Baker et al. 2018). Additionally, environmental conditions (e.g., chlorophyll α, natural inhibitors, microbial activity, biochemical oxygen demand (BOD), temperature, pH, and ultraviolet B (UV-B) radiation) play an integral role in eDNA degradation rates (Barnes et al. 2014; Pilliod et al. 2014; Strickler et al. 2015; Lance et al. 2017; Stoeckle et al. 2017; Seymour et al. 2018).

The complex nature of eDNA has led to a new branch of eDNA research that aims to disentangle the factors influencing its ecology, such as the distribution of eDNA across both spatial and temporal scales (Spear et al. 2015; Wilcox et al. 2016; Tillotson et al. 2018), and a mechanistic understanding of eDNA ecology in relation to transport, retention (i.e., deposition or capture by sediment) and subsequent resuspension (Jane et al. 2015; Jerde et al. 2016; Shogren et al. 2016; Shogren et al. 2017).

The majority of the aforementioned studies have targeted single species using real-time quantitative PCR (qPCR) or droplet digital PCR (ddPCR) to investigate eDNA ecology. Recently, eDNA metabarcoding, which combines PCR amplification with High-Throughput Sequencing (HTS), has emerged as a powerful, efficient, and economical tool for biodiversity assessment and monitoring of entire aquatic communities (e.g., Deiner et al. 2016; Hänfling et al. 2016; Port et al. 2016; Valentini et al. 2016). This tool removes the need to select target organisms *a priori* with the use of generic PCR primers that amplify multiple taxa, thus facilitating detection of invasive or threatened species when conducting holistic biodiversity assessment and routine freshwater monitoring (Thomsen & Willerslev 2015). Encouragingly, Harper et al. (2018) demonstrated that *Triturus cristatus* (great crested newt) detection via metabarcoding with no threshold is equivalent to qPCR with a stringent detection threshold. eDNA metabarcoding has also been applied to large-scale investigations of spatial or temporal variation in marine and freshwater communities, with some studies indicating that communities can be distinguished from 100 m to 2 km due to stream discharge or tidal patterns (Civade et al. 2016; Port et al. 2016; O’Donnell et al. 2017; Kelly et al. 2018; Li et al. 2018b).

In this study, we capitalise on the diagnostic power of eDNA metabarcoding to explore the spatial and temporal distribution of fish communities in two aquaculture ponds and evaluate the detection sensitivity of this tool for low-density species alongside highly abundant species. Two primary objectives are investigated. Firstly, the shedding and decay rates of eDNA in fish ponds are explored, following the introduction and removal of two rare species at a fixed location. Secondly, the spatial distribution of fish communities after rare species introduction and removal is examined. We expect that eDNA would be shed and diffused away from its source (the rare and introduced species), and this increased movement of eDNA particles would homogenise β-diversity in terms of community similarity, thus eroding the distance-decay relationship of eDNA. The results of this research are critical for understanding the characteristics of eDNA in ponds including production, degradation and transport, and to inform effective sampling strategies.

## 2 Materials and Methods

### 2.1 Study site and water sampling

This study was carried out at two artificially stocked ponds with a high fish density and in a turbid and eutrophic condition. The two ponds (E1 and E4) are located at the National Coarse Fish Rearing Unit (NCFRU, Calverton, Nottingham, UK), run by the UK Environment Agency. The ponds are groundwater fed with no inflow from surface water bodies. The dimension of each pond is approximately 60 m × 85 m, with an average depth of 1.5 m. In each pond, there were two feeding devices with timers that release food hourly, and two automatic aerators near the feeding devices to increase the dissolved oxygen (DO) profile. The automatic aerators also created flowing conditions for the fish to feed in and to help build the right kind of muscle needed for life in the wild (Fig. 1). Generally, these ponds are used to rear approximately one-year-old common British coarse fish before they are used in stocking programmes for conservation purposes or recreational fishing.

**Figure 1.**
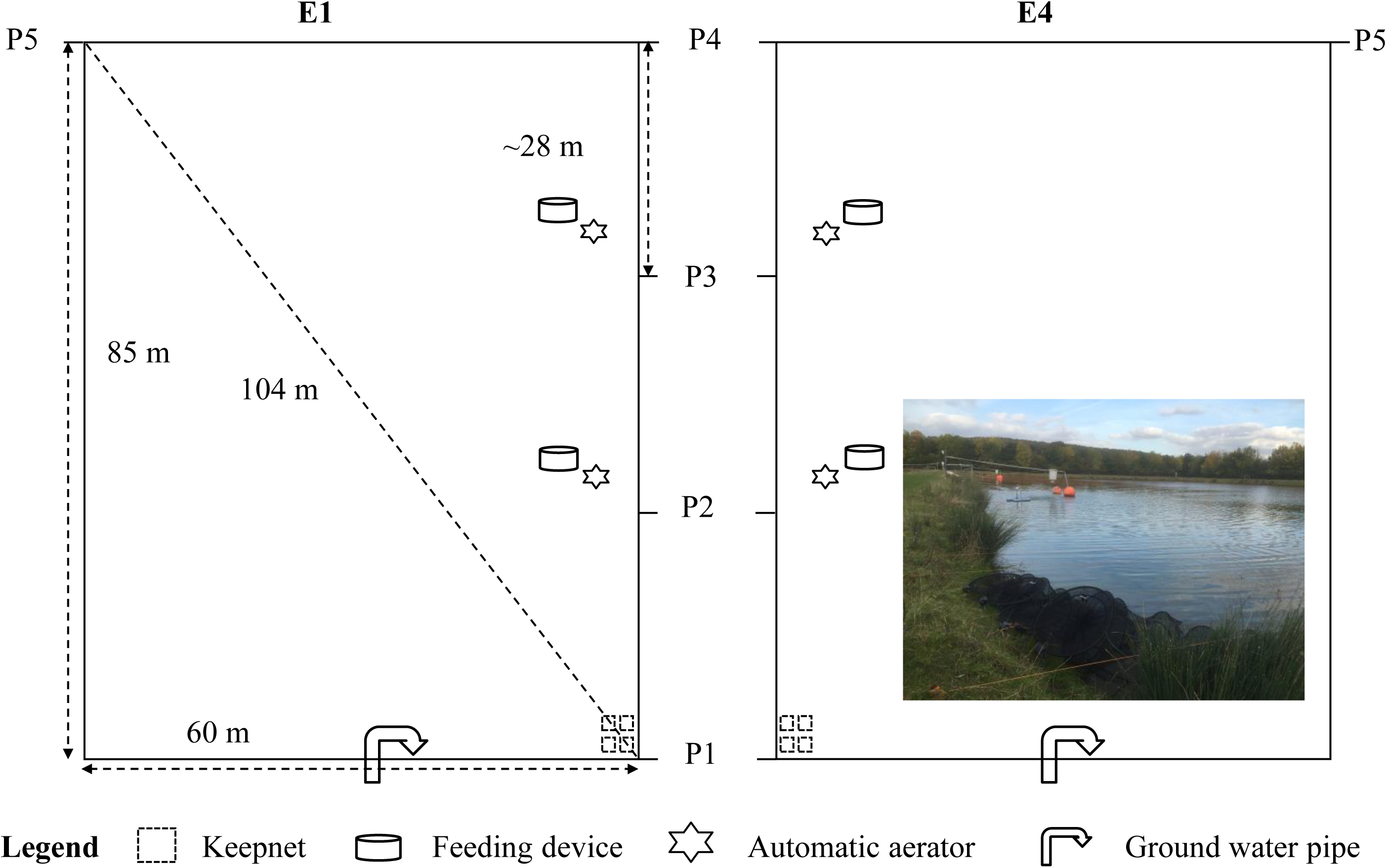
Schematic of sampling strategy at the National Coarse Fish Rearing Unit. The linear distance of each sampling position to keepnets with the introduced species is 0 m (P1), 28 m (P2), 56 m (P3), 84 m (P4), and 104 m (P5).

The experiment was conducted from 19^th^ September to 3^rd^ October 2016. DO and temperature were monitored daily in each pond during the entire sampling period. DO concentration and temperature were 8.4 ± 1.3 mg L^-1^ and 15.6 ± 1.4 °C in pond E1, and 7.1 ± 1.4 mg L^-1^ and 16.0 ± 1.3 °C in pond E4. Stocked fish in both ponds were measured and weighed before stocking on 16^th^ June 2016 and after harvesting on 18^th^ November 2016. Fish abundance and biomass at time of water sampling in September 2016 were estimated, assuming that the death and growth curves of these fish are linear (Supporting Information Figs. A1 & A2). The fish stock information in September 2016 is shown in Table 1. On 19^th^ September at 15:00 (hereafter referred to as “D0”), an hour prior to introduction of additional fish species, one 2 L water sample was taken just below the pond surface using sterile Gosselin™ HDPE plastic bottles (Fisher Scientific) at each of the five sampling positions (hereafter referred to as “P1–P5”) spread over 104 m, to confirm fish community composition and check for potential contamination from aberrant species. Briefly, four sampling positions (P1–P4) were distributed equidistant on the same shoreline of the pond, whereas P5 was on the catercorner of P1 (Fig. 1). After sampling on D0, four new keepnets containing 25 individuals each of the introduced species were placed in P1 of each pond. In pond E1, the introduced species were *Squalius cephalus* (chub, 26.0 ± 1.8 g) and *Scardinius erythrophthalmus* (rudd, 21.8 ± 1.5 g), whereas rudd (22.4 ± 1.6 g) and *Leuciscus leuciscus* (dace, 19.8 ± 1.5 g) were introduced to pond E4. After fish introduction, five 2 L water samples were collected at 10:00 on days 2, 4, 6 and 8 (hereafter referred to as “D2–D8”, introductory stage) at each position (P1–P5) in each pond. On D8, the keepnets with introduced species were removed after water sampling on that day was completed. No fish died in the keepnets. After removal of the keepnets, water samples were collected in the same manner on days 10, 12 and 14 (hereafter referred to as “D10–D14”, removal stage) in order to estimate eDNA decay of the introduced species once removed from the pond. In each pond, forty samples were taken over the course of the experiment (80 samples in total). The introduced species were weighed after removal from ponds, and then released back into indoor tanks at NCFRU. All animal research was approved by the University of Hull’s Faculty of Science Ethics Committee (Approval #U093).

**Table 1.** Fish stock information on two experimental ponds at the National Coarse Fish Rearing Unit.

| Pond | Species |  |  | September 2016 |  |
| --- | --- | --- | --- | --- | --- |
|  | Scientific name | Common name | Code | Abundance | Biomass (kg) |
| E1 | *Barbus barbus* | Barbel | BAR | 7245 | 267.99 |
| E1 | *Abramis brama* | Bream | BRE | 6449 | 152.33 |
| E1 | *Carassius carassius* | Crucian carp | CAR | 2309 | 80.44 |
| E1 | *Squalius cephalus*† | Chub | CHU | 50 | 1.30 |
| E1 | *Leuciscus leuciscus* | Dace | DAC | 18544 | 123.96 |
| E1 | *Rutilus rutilus* | Roach | ROA | 3452 | 44.64 |
| E1 | *Scardinius erythrophthalmus*† | Rudd | RUD | 50 | 1.09 |
| E1 | *Tinca tinca* | Tench | TEN | 3605 | 59.09 |
| E4 | *Barbus barbus* | Barbel | BAR | 4230 | 165.07 |
| E4 | *Abramis brama* | Bream | BRE | 1130 | 32.33 |
| E4 | *Carassius carassius* | Crucian carp | CAR | 1766 | 79.25 |
| E4 | *Squalius cephalus* | Chub | CHU | 16395 | 492.01 |
| E4 | *Leuciscus leuciscus*† | Dace | DAC | 50 | 0.99 |
| E4 | *Rutilus rutilus* | Roach | ROA | 24732 | 355.53 |
| E4 | *Scardinius erythrophthalmus*† | Rudd | RUD | 50 | 1.12 |
| E4 | *Tinca tinca* | Tench | TEN | 645 | 9.28 |

Notes: “†” indicates the rare species introduced to each pond for the purposes of this study.
Abundance represents number of individuals. Full scientific, common names and three letter codes used in figures and tables are given.

### 2.2 eDNA capture and extraction

After each sampling event, all water samples were filtered immediately in a laboratory at NCFRU that was decontaminated before filtration by bleaching (50% v/v commercial bleach) floors and surfaces. Three filtration replicates (300 mL) were subsampled from each 2 L water sample collected at every sampling position. All filtration replicates were filtered through sterile 0.8 μm mixed cellulose acetate and nitrate (MCE) filters, 47 mm diameter (Whatman) using Nalgene filtration units in combination with a vacuum pump (15–20 in. Hg; Pall Corporation). Our previous study demonstrated that 0.8 μm is the optimal membrane filter pore size for turbid, eutrophic, and high fish density ponds, and achieves a good balance between rapid filtration time and the probability of species detection via metabarcoding (Li et al. 2018a).

To reduce cross-contamination, samples from the same pond were filtered in the same batch and in order of collection from P1 to P5. The same filtration unit was used for all three filtration replicates of each sample. The filtration units were soaked in 10% v/v commercial bleach solution 10 mins and 5% v/v microsol detergent (Anachem, UK) 5 mins, and then rinsed thoroughly with deionised water after each round of filtration to prevent cross-contamination. One filtration blank (300 mL deionised water) was processed for each pond on every day of filtration to monitor contamination risk. After filtration, all membrane filters were placed into 50 mm sterile petri dishes (Fisher Scientific) using sterile tweezers, sealed with Parafilm^®^ (Bemis Company, Inc.), and stored at –20 °C until DNA extraction. DNA extraction was carried out using the PowerWater^®^ DNA Isolation Kit (MoBio Laboratories Inc., now QIAGEN) following the manufacturer’s protocol. The DNA was eluted in 100 μL 10 mM Tris (Solution PW6) and stored at –20 °C freezer.

### 2.3 Library preparation and sequencing

Extracted DNA samples were amplified with vertebrate-specific primers (Riaz et al. 2011) that target a 106-bp fragment of the mitochondrial 12S rRNA region in fish, using a two-step PCR protocol for library preparation that implements a nested tagging approach (Kitson et al. 2018). Previous eDNA metabarcoding studies of marine mesocosms and coastal ecosystems showed that this fragment has a low false negative rate for bony fishes (Kelly et al. 2014; Port et al. 2016). We also previously tested this fragment *in situ* on three deep lakes in the Lake District, England, where metabarcoding results were compared to long-term data from established survey methods (Hänfling et al. 2016), and at NCFRU to investigate the impact of different filters on eDNA capture and quantification (Li et al. 2018a). Taken together, our previous findings demonstrated that this 106-bp fragment is highly suitable for eDNA metabarcoding of UK freshwater fish communities.

In the two-step library preparation protocol, the first PCR reactions were set up in a UV and bleach sterilised laminar flow hood in our dedicated eDNA laboratory at the University of Hull to minimise contamination risk. All filtration replicates (*N* = 240), together with 16 filtration and extraction blanks, 16 no-template controls (NTCs), and 16 single-template positive controls (STCs) were included in library construction (*N* = 288) for sequencing on an Illumina MiSeq. For the STCs, we used genomic DNA (0.08 ng uL^-1^) of *Astatotilapia calliptera* (Eastern happy), a cichlid from Lake Malawi that is not present in natural waters in UK.

The first PCR reaction was carried out in 25 μL volumes containing: 12.5 μL of 2 × MyTaq HS Red Mix (Bioline), 0.5 μM of each tagged primer, 2.5 μL of template DNA, and 7.5 μL of molecular grade water. Eight-strip PCR tubes with individually attached lids and mineral oil (Sigma-Aldrich) were used to reduce cross-contamination between samples. After PCR preparation, reaction tubes were brought to our PCR room for amplification, where all post-PCR work was carried out. Thermal cycling parameters were as follows: 98 °C for 5 min, 35 cycles of 98 °C for 10 sec, 58 °C for 20 sec, and 72 °C for 30 sec, followed by a final elongation step at 72 °C for 7 min. Three PCR technical replicates were performed for each sample, then pooled to minimise bias in individual PCRs. The indexed first PCR products of each sample were then pooled according to sampling event and pond, and 100 μL of pooled products cleaned using the Mag-Bind^®^ RXNPure Plus Kit (Omega Bio-tek) using a dual bead-based size selection protocol (Bronner et al. 2014). Ratios used for size selection were 0.9× and 0.15× magnetic beads to PCR product.

The second PCR reactions were carried out in 50 μL volumes containing: 25 μL 2 × MyTaq HS Red Mix (Bioline), 1.0 μM of each tagged primer, 5 μL of template DNA and 15 μL of molecular grade water. Reactions without template DNA were prepared in our dedicated eDNA laboratory, and first PCR products added later in the PCR room. Thermal cycling parameters were as follows: initial denaturation at 95 °C for 3 min, followed by 10 cycles of 98 °C for 20 s, and 72 °C 1 min, with a final extension of 72 °C for 5 min. The second PCR products (50 μL) were cleaned using the Mag-Bind^®^ RXNPure Plus Kit (Omega Bio-tek) according to a dual bead-based size selection protocol (Bronner et al. 2014). Ratios used for size selection were 0.7× and 0.15× magnetic beads to PCR product. The cleaned second PCR products were normalised according to sample number and concentration across sampling events and ponds based on the Qubit™ 3.0 fluorometer results using a Qubit™ dsDNA HS Assay Kit (Invitrogen), then pooled. The final library concentration was quantified by qPCR using the NEBNext^^®^^ Library Quant Kit (New England Biolabs). The pooled, quantified library was adjusted to 4 nM and denatured following the Illumina MiSeq library denaturation and dilution guide. To improve clustering during initial sequencing, the denatured library (13 pM) was mixed with 10% PhiX genomic control. The library was sequenced on an Illumina MiSeq platform using the MiSeq reagent kit v2 (2 × 250 cycles) at the University of Hull.

### 2.4 Data analysis

#### 2.4.1 Bioinformatics analysis

Raw read data from the Illumina MiSeq have been submitted to NCBI (BioProject: PRJNA486650; BioSample accession: SAMN09859568–SAMN09859583; Sequence Read Archive accessions: SRR7716776–SRR7716791). Bioinformatics analysis was implemented using a custom, reproducible pipeline for metabarcoding data (metaBEAT v0.97.10) with a custom 12S UK freshwater fish reference database (Hänfling et al. 2016). Sequences for which the best BLAST hit had a bit score below 80 or had less than 100% identity to any sequence in the curated database were considered non-target sequences. To assure full reproducibility of our bioinformatics analysis, the custom 12S reference database and the Jupyter notebook for data processing have been deposited in a dedicated GitHub repository (https://github.com/HullUni-bioinformatics/Li_et_al_2018_eDNA_dynamic). The Jupyter notebook also performs demultiplexing of the indexed barcodes added in the first PCR reactions.

#### 2.4.2 Criteria for reducing false positives and quality control

Filtered data were summarised as the number of sequence reads per species (hereon referred to as read counts) for downstream analyses (Supporting Information Appendix S1). After bioinformatics analysis, the low-frequency noise threshold (proportion of STC species read counts in the real sample) was set to 0.002 to filter out high-quality annotated reads that passed the previous filtering steps and had high-confidence BLAST matches, but may have resulted from contamination during the library construction process or sequencing (De Barba et al. 2014; Hänfling et al. 2016; Port et al. 2016). The low-frequency noise threshold was guided by the analysis of sequence data from STCs under several thresholds.

#### 2.4.3 Statistical and ecological analyses

All statistical analyses were performed in R v3.5.0 (R_Core_Team 2018), and graphs were plotted using GGPLOT2 v2.2.1 (Wickham & Chang 2016). The sequence read counts of different filtration replicates (*N* = 3) were averaged to provide a single read count for each sampling position unless otherwise specified. The fish community of each sampling position was standardised to proportional abundance (i.e., number of read counts per species relative to total number of read counts in that sample) using the “total” method with the function *decostand* in VEGAN v2.4-4 (Oksanen et al. 2017). To evaluate spatial and temporal species turnover between eDNA communities, the observed variation in distance measured as Bray-Curtis dissimilarity among sampling events and positions were apportioned using permutational multivariate analysis of variance (PERMANOVA) with the function *adonis* in VEGAN v2.4-4 (Oksanen et al. 2017). To determine the relationship between β-diversity in Bray-Curtis distance matrices of different sampling days (D0–D14) and the geographic distance matrix of different sampling positions (P1–P5), the Mantel correlations were performed with the function *mantel.rtest* of ADE4 v1.7-11 (Stéphane et al. 2018). To examine temporal and spatial variance in fish communities after the introduction and removal of introduced species, pairwise Bray-Curtis dissimilarities were calculated using the function *vegdist* in VEGAN v2.4-4 (Oksanen et al. 2017), and Kruskal-Wallis one-way ANOVA with Dunn’s test using Bonferroni adjustment conducted to test for differences in Bray-Curtis dissimilarity between different sampling days and positions. The statistical significance level of this study is set at 0.05. The full R script is available on the GitHub repository (https://github.com/HullUni-bioinformatics/Lietal2018eDNAdynamic/tree/master/Rscript).

## 3 Results

The library generated 16.99 million reads with 13.21 million reads passing filter including 10.94% PhiX control. Following quality filtering and removal of chimeric sequences, the average read count per sample (excluding controls) was 14,441. After BLAST searches for taxonomic assignment, 51.50% ± 10.87% reads in each sample were assigned to fish.

### 3.1 Species detection in the background communities

All stocked species were detected over the course of the experiment in ponds E1 and E4. In pond E1, the stocked species were *Abramis brama* (common bream), *Barbus barbus* (barbel), *Carassius carassius* (crucian carp), dace, *Rutilus rutilus* (roach) and *Tinca tinca* (tench). In pond E4, the stocked species were common bream, barbel, crucian carp, chub, roach, and tench (Fig. 2). Moreover, apart from tench in pond E4, stocked species were detected across all sampling positions (Fig. 2; Supporting Information Table A1). Tench was the rarest stocked species in pond E4 (proportional individual and biomass was 1.32% and 0.82%, respectively; Fig. 2B; Table 1) which may explain imperfect species detection.

**Figure 2.**
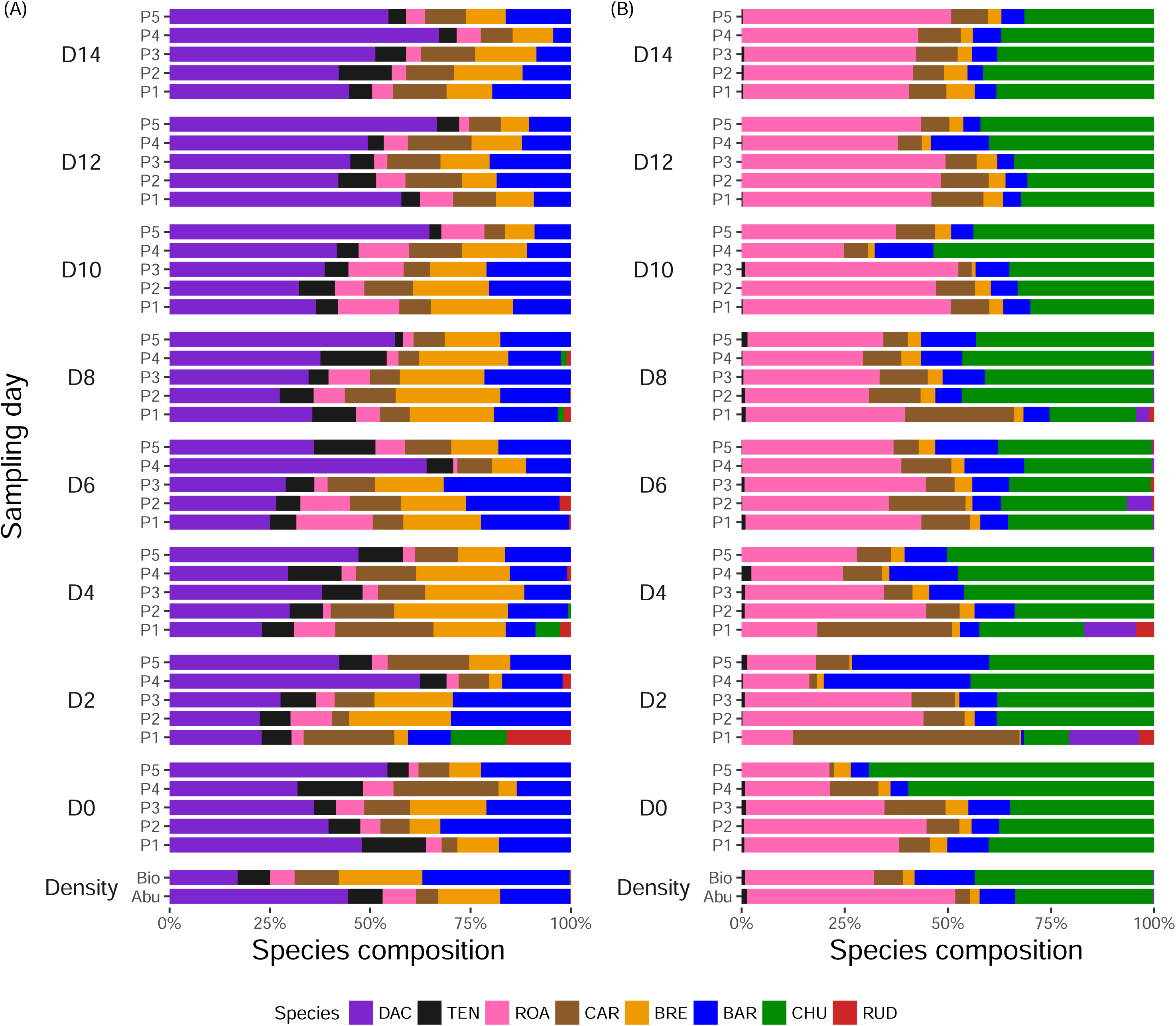
Species composition of averaged read counts (number of replicates = 3) for five sampling positions over 14 days in ponds (A) E1 and (B) E4. “Bio” and “Abu” refers to fish biomass and abundance density respectively, calculated based on Table 1. Species three letter codes correspond to species given in Table 1. After control samples were taken on D0, the rare species were introduced and samples were taken on days 2, 4, 6, 8, 10, 12, and 14 (D2–D14) from the five sampling positions (P1–P5). The introduced species were removed on D8 after sampling. The linear distance of each sampling position to keepnets of introduced species is 0 m (P1), 28 m (P2), 56 m (P3), 84 m (P4), and 104 m (P5).

### 3.2 Spatio-temporal detection of introduced species

The introduced species were not detected in samples taken prior to species introduction (i.e., D0), or in process controls (filtration, extraction and NTCs). Therefore, the introduced species were not present in the environment or as laboratory contaminants before the experiment began. After introduction of rudd and chub into pond E1, rudd were detected across the entire period the species were present (D2–D8), whereas chub were not recovered on D6 in pond E1. In pond E4, both the introduced species, rudd and dace, were identified across the entire period the species were present (Fig. 2). In terms of sampling position, the eDNA signal of the introduced species was strongest close to the keepnets (P1) and decreased with increasing distance from this location (Fig. 2). In pond E1, both introduced species were detected until P4 (85 m from the keepnets), but not at the catercorner of the keepnets (P5, 104 m away from the keepnets). In contrast, in pond E4, both introduced species could be detected at P5 on D6 (Fig. 3). The detection probability of the introduced species at P1 across both ponds (Supporting Information Table A2, 0.88 ± 0.13) was significantly higher than other sampling positions during the entire period the species were present (Supporting Information Table A2, ANOVA, *p* consistently < 0.05). Moreover, eDNA concentration (i.e., proportional read counts abundance) of introduced species was highest on D2 at the original source (P1) in both ponds (Fig. 3A, F). Thereafter, eDNA concentration decreased gradually and reached equilibrium (i.e., the production rate equal to degradation rate) on D6, with a slight increase on D8 (Fig. 3A, F). There was also some variation in eDNA concentration among species that was unrelated to fish density. For instance, the eDNA concentration of rudd was higher than chub in pond E1 but lower than dace in pond E4 (Fig. 3), even though the biomass of rudd was lower than chub in pond E1 and higher than dace in pond E4 (Table 1). Notably, after the introduced species had been removed for 48 hrs (D8–D10), they were no longer detectable at any position in both ponds (Figs. 2 & 3).

**Figure 3.**
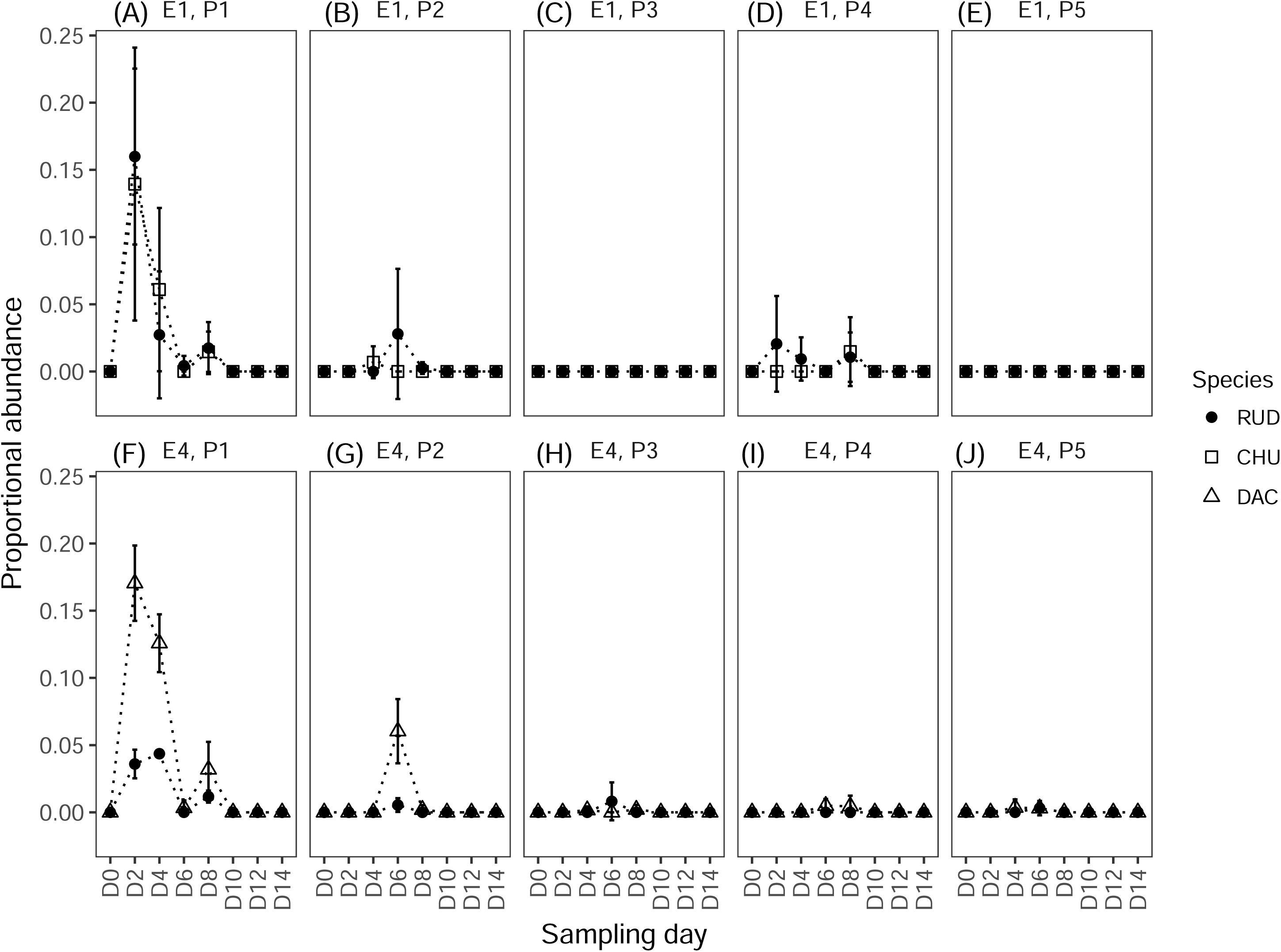
Temporal change over 14 days (D0–D14) in averaged proportional abundance of introduced species across five sampling positions (P1–P5) in ponds E1 and E4. The standard error bars represent three filtration replicates per sample. Species three letter codes correspond to species given in Table 1. The different sampling stages and linear distance between sampling positions are described in Fig. 2.

### 3.3 Community variance in Bray-Curtis dissimilarity

On the whole, sampling day and position had significant effects on community variance, using Bray-Curtis dissimilarity for ponds E1 (PERMANOVA; sampling days *df* = 7, *R^2^* = 0.296, *p* = 0.002; positions *df* = 4, *R^2^* = 0.235, *p* = 0.002) and E4 (PERMANOVA; sampling days *df* = 7, *R^2^* = 0.241, *p* = 0.013; positions *df* = 4, *R^2^* = 0.271, *p* = 0.001). Specifically, the estimates of community dissimilarity for different sampling positions between different sampling days were not correlated with geographic distance, except D0 in pond E4. Moreover, there were significant correlations of community dissimilarity between D8, D10 and D12, D6 and D14 in pond E1. Significant correlations of community dissimilarity were observed between D0 and D14, D2 and D4, D2 and D8, D10 and D12 in pond E4. All the *r* statistics and *p*-values as determined by the Mantel test are shown in Fig. 4.

**Figure 4.**
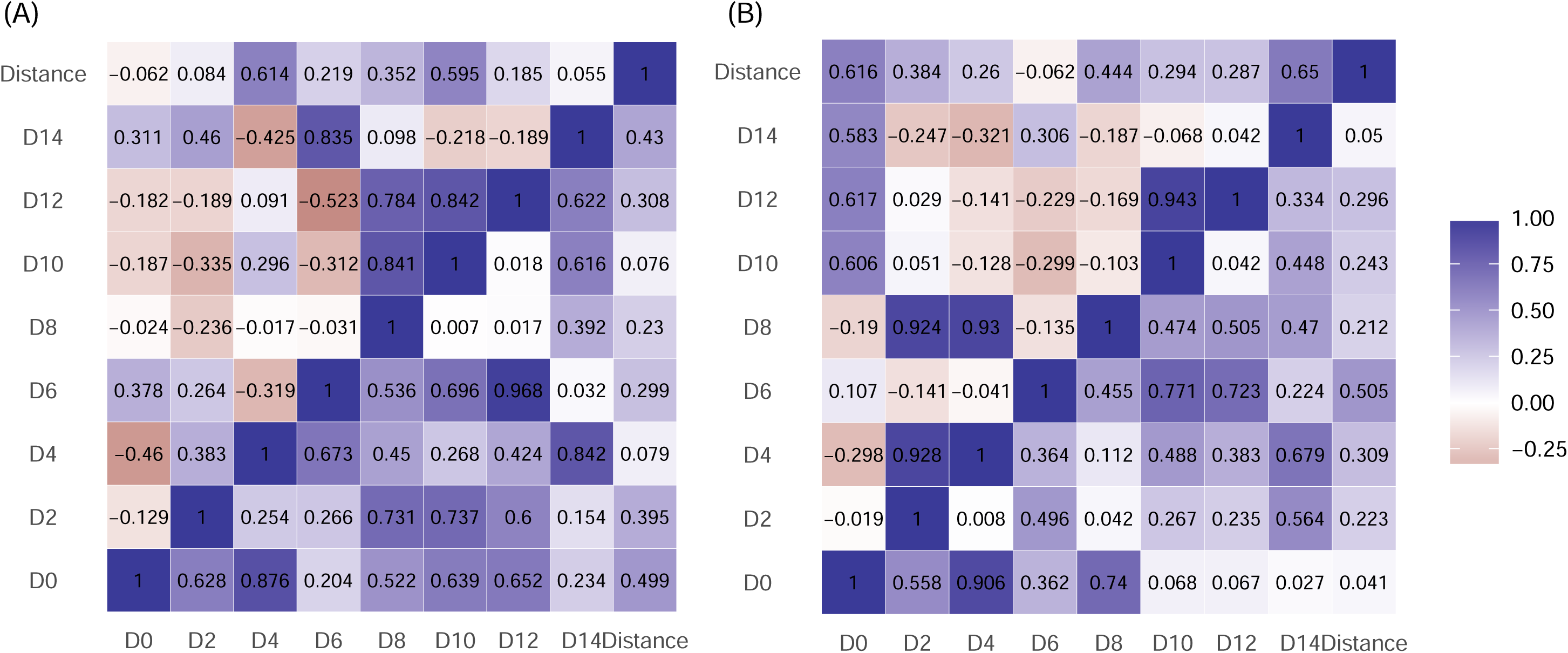
Heatmap of community correlation as determined by the Mantel test between Bray-Curtis distance matrices of different sampling days (D0–D14) and the geographic distance matrix of different sampling positions (P1–P5) in ponds (A) E1 and (B) E4. “Distance” refers to the distance matrix based on the linear distance between different sampling positions. The upper triangular and lower triangular is Mantel *r* statistics and *p*-values, respectively. The different sampling stages and linear distance between sampling positions are described in Fig. 2.

Overall, fish communities varied in Bray-Curtis dissimilarity before introduction on D0, introduction from D2 to D8, and removal from D10 to D14 (Supporting Information Fig. A3). The Bray-Curtis dissimilarity of the removal stage was significantly lower than the introductory stage in both ponds E1 and E4 (Supporting Information Fig. A3; Dunn’s test; E1 z = 3.71, *p* < 0.05; E4 z = 2.98, *p* < 0.05). In pond E4, community dissimilarity of the removal stage was also significantly lower than before the introduction of species (Supporting Information Fig. A3B; Dunn’s test, z = 2.45, *p* < 0.05). More specifically, after the introduction of rare species, the highest community dissimilarities of different sampling positions were observed on D2 and decreased over time in both ponds. There was no significant difference between sampling days during D4–D14 and D6–D14 in ponds E1 and E4, respectively (Fig. 5A1, B1). In terms of sampling position, the highest community variances of different sampling days occurred close to the keepnets (P1) in both ponds (Fig. 5A2, B2), and the community dissimilarity significantly declined with increasing distance from P1 to P3. However, communities were more dissimilar at P4 compared to P3, with a significant increase in Bray-Curtis dissimilarity values (Fig. 5A2, B2; Dunn’s test, E1 z = 2.92, *p* < 0.05, E4 z = 2.95, *p* < 0.05). In pond E1, there was a significant reduction in community dissimilarity at P5 compared to P4 (Fig. 5A2; Dunn’s test, z = 2.83, *p* < 0.05), whereas in pond E4, there was no significant difference in community dissimilarity between P4 and P5 (Figure 5b2).

**Figure 5.**
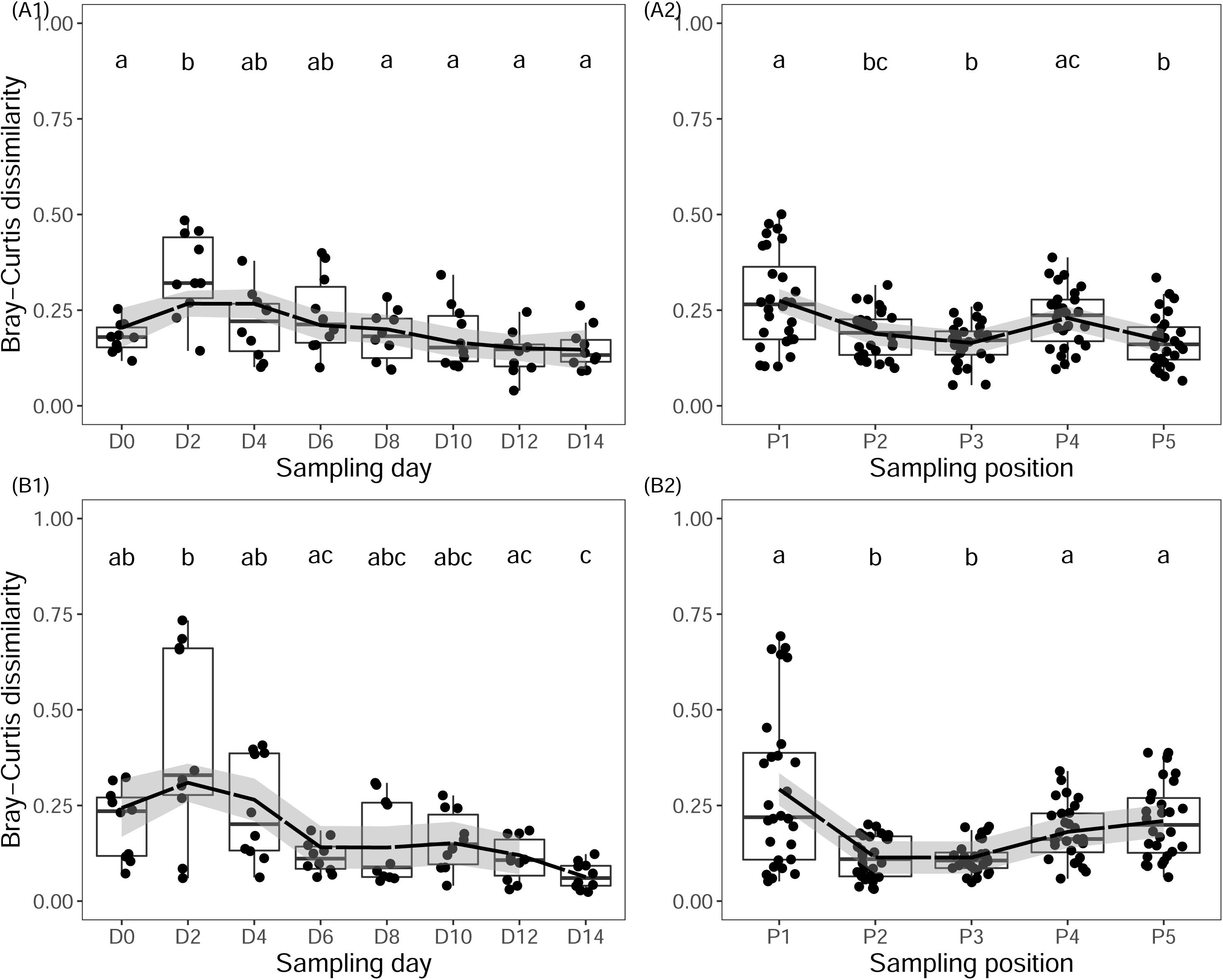
Temporal change (D0–D14) in community dissimilarity of the five sampling positions (P1–P5) in ponds (A1) E1 and (B1) E4, where each point represents the Bray-Curtis dissimilarity of two different sampling positions on the same sampling day. Spatial change (P1–P5) in community dissimilarity of the eight sampling days (D0–D14) in ponds (A2) E1 and (B2) E4, where each point represents the Bray-Curtis dissimilarity of two different sampling days at the same sampling position. Sampling days or positions that differ significantly (*p* < 0.05) from one another are indicated with different letters in each boxplot. Dashed lines represent the fit of non-linear regressions, and grey shaded areas denote the 95% confidence interval as calculated using the standard error. The different sampling stages and linear distance between sampling positions are described in Fig. 2.

## 4 Discussion

Spatial heterogeneity of eDNA distribution has been reported in lentic ecosystems (Takahara et al. 2012; Eichmiller et al. 2014; Hänfling et al. 2016; Lawson Handley et al. 2018). Therefore, an understanding of the spatial heterogeneity of eDNA distribution is critical to the design of effective sampling protocols for accurate species detection and abundance estimates in lentic ecosystems, especially in order to detect rare or invasive species. To our knowledge, this study is the first that uses metabarcoding to investigate the spatial and temporal community variances in ponds to understand eDNA ecology in these systems, including production, degradation and transport following the introduction and removal of rare species.

### 4.1 eDNA production

The eDNA concentration of the introduced species peaks on D2 at the position closest to the keepnets (P1). Thereafter, eDNA concentration of these introduced species declines gradually over time and stabilises by D6 in both ponds. Consequently, the highest community dissimilarity of different sampling positions is observed on D2 and decreases over time in both ponds. The increase in eDNA concentration of the introduced fish after 43 hrs may have been caused by increased eDNA shedding rates as a result of fish being stressed by handling, as observed in other studies (Takahara et al. 2012; Maruyama et al. 2014; Klymus et al. 2015; Sassoubre et al. 2016). Considering the degradation rate of eDNA is less than 48 hrs (see more detail in Section 4.2), eDNA concentration may have declined after D2 due to fish acclimation to the keepnets and reduced activity, resulting in less eDNA release. By D6, the rate of eDNA release from the two introduced species seems to reach equilibrium with the rate of eDNA degradation. These patterns are consistent with previous qPCR studies that targeted single species and investigated eDNA production and degradation, including eDNA shedding rate of *Cyprinus carpio* (common carp) in aquaria (Takahara et al. 2012), different developmental stages of *Lepomis macrochirus* (bluegill sunfish) in aquaria (Maruyama et al. 2014) and three marine fish, *Engraulis mordax* (Northern anchovy), *Sardinops sagax* (Pacific sardine) and *Scomber japonicas* (Pacific chub mackerel), in seawater mesocosms (Sassoubre et al. 2016). However, eDNA concentration of two amphibian species, *Pelobates fuscus* (common spadefoot toad) and great crested newt, exhibit monotonic increases after introduction into aquaria, which may be the result of a longer sampling period over larger time intervals, i.e., weeks over two months or lower degradation rates in controlled environments (Thomsen et al. 2012b).

### 4.2 eDNA degradation

The detection rates of the introduced species declines with no detectable eDNA signal at any sampling position in both ponds approximately 48 hrs after removal. As a result, there is no significant difference in community dissimilarity of different sampling positions among the sampling days after removal of the introduced species. This observation is in agreement with other studies that documented no eDNA detection shortly after target species were removed from the water in which they occurred. For example, detection of *Platichthys flesus* (European flounder) or bluegill sunfish fails around 24 hours after removal from aquaria (Thomsen et al. 2012a; Maruyama et al. 2014), and 48 hrs after removal of *Salmo salar* (Atlantic salmon) from a river ecosystem (Balasingham et al. 2017). By contrast, other studies have reported slower eDNA degradation rates in controlled aquaria or mesocosms. For example, eDNA degrades beyond detection within a week for fish (Thomsen et al. 2012a; Barnes et al. 2014; Sassoubre et al. 2016), several weeks for amphibians (Dejean et al. 2011; Thomsen et al. 2012b), and a month for *Potamopyrgus antipodarum* (New Zealand mudsnail) (Goldberg et al. 2013), The wide variation observed in the aforementioned studies emphasises the role of the ecosystem and starting eDNA concentration (influenced by shedding rate) on eDNA persistence. The reason for wide variation in eDNA production rates among species is unconfirmed, but animal physiology is suggested to play a role, e.g., stress (Pilliod et al. 2014), breeding readiness (Spear et al. 2015), diet (Klymus et al. 2015), and metabolic rate (Maruyama et al. 2014). Moreover, eDNA is also found to decay faster in the field than in controlled conditions, which can be attributed to the complex effects of environmental conditions on eDNA persistence (Barnes et al. 2014; Pilliod et al. 2014; Strickler et al. 2015; Lance et al. 2017; Stoeckle et al. 2017; Seymour et al. 2018).

### 4.3 eDNA transport

Regarding horizontal transport of eDNA, the eDNA signal and detection probability of the introduced species is highest close to the keepnets (P1) and broadly decreases with increasing distance up to around 104 m from this point. This finding agrees with previous qPCR studies that reported a patchy distribution of eDNA in the lentic ecosystems, and drastic decline in detection probability and eDNA concentration less than a few hundred metres from the target organisms (Takahara et al. 2012; Eichmiller et al. 2014; Dunker et al. 2016). Moreover, all estimates of β-diversity (i.e., community dissimilarity) of different sampling positions between different sampling days and geographic distances are not linearly correlated, except D0 in pond E4, which indicates that geographic distance does not have a significant effect. This result would imply that the eDNA of stocked fish is well homogenised in the ponds, and the eDNA signal released by the introduced species is too low to influence the spatial distribution pattern of the entire fish community present in the ponds. This result is in agreement with Evans et al. (2017) who do not find a significant relationship between sample dissimilarity and geographic distance in a 22,000 m^2^ surface-area reservoir in which fish distribution is relatively homogeneous. By contrast, Sato et al. (2017) indicated that geographic distances among sampling locations within lakes ranging in size from 84,000 m^2^ to 2,219,000 m^2^ have a significantly positive correlation with the abundance-based community dissimilarity index resulting from spatial heterogeneity of eDNA distribution.

In lotic ecosystems, stream discharge plays an important role in horizontal eDNA transport, and can result in eDNA of target species being transported metres to kilometres (Deiner & Altermatt 2014; Pilliod et al. 2014; Jane et al. 2015; Jerde et al. 2016). Furthermore, the spatial community variance observed in other eDNA studies indicated that β-diversity does not increase as a function of distance (up to 12 km) in a stream (Deiner et al. 2016), but does increase with distance in a highly dynamic marine habitat (O’Donnell et al. 2017). Li et al. (2018b) also observed that the β-diversity of fish communities based on Jaccard distance (i.e., incidence data) between sampling sites is correlated with the sampling distance along the stream.

In the small fish ponds sampled in this study, the community variance in eDNA distribution is highly localised in space. The cline of community variance over distance is consistent, where eDNA signal of the introduced species is strongest at the position closest to the keepnets (P1), followed by a reduction in strength from P1 to P3 and growth from P3 to P4. Furthermore, two introduced species are detected at P5 in pond E4, but not at P5 in pond E1. This may explain why there is no significant change in community dissimilarity between P4 and P5 in pond E4, but the community dissimilarity of P5 is significantly reduced from P4 in pond E1. Notably, there are feeding devices and automatic aerators near P2 and P3. Thus, we speculated that food released by feeding devices could attract fish and cause them to aggregate near positions P2 and P3, which would increase the detection of stocked fish and thus reduce the detection probabilities of the introduced species. On the other hand, the automatic aerators could have enhanced water mixing, bringing eDNA from the introduced species into the other corner of the pond (P4). Therefore, the growth trend in eDNA concentration of the introduced species from P3 to P4 in both ponds may be a consequence of anthropogenic interference.

## 5 Conclusions

This study has demonstrated that eDNA metabarcoding is a powerful tool for monitoring change in community structure across time and space. After eDNA is shed and transported away from its source, the increased movement of eDNA particles homogenises community similarity and erodes the distance-decay relationship of eDNA. Notably, after two introduced species have been removed, they are not detectable at any sampling position after 48 hrs. These findings on the spatial and temporal resolution of eDNA support that genetic material present in static environments originates from organisms that are nearby or have been nearby very recently. This work serves as an important case study of eDNA-based community diversity at fine temporal and spatial scales in ponds as a coherent view of eDNA ecology and dynamics begins to come into focus. While our observations are instructive, further quantitative modelling of eDNA transport, retention, and subsequent resuspension are needed to predict species location and estimate abundance (e.g., Jane et al. 2015; Jerde et al. 2016; Shogren et al. 2016; Shogren et al. 2017). This will be critical to take eDNA analysis to the next level as a powerful, diagnostic tool in ecology, conservation, and management. Regardless of modelling approaches, rigorous and spatially standardised sampling designs are key to ensuring the reliability of eDNA surveillance.

## Acknowledgments

This work is part of the PhD project of J.L., who is supported by the University of Hull and the China Scholarship Council. We are particularly grateful to all staff (Alan Henshaw, Richard Pitman, Nick Gill, Dan Clark and James Rabjohns) of the National Coarse Fish Rearing Unit of the UK Environment Agency for their help with sampling and providing fish stock information, and Drs Christoph Hahn, Amir Szitenberg, Peter Shum and Rob Donnelly for their assistance with bioinformatics analysis and lab work.

## Data accessibility

Data is available from the GitHub repository https://github.com/HullUni-bioinformatics/Li_et_al_2018_eDNA_dynamic). The repository will be archived with Zenodo after the manuscript accepted.

## Author contributions

The study was conceived and designed by B.H. and J.L.; sampling were conducted by J.L. L.R.H., and H.V.W; experiment and bioinformatics analyses were performed by J.L.; statistical analyses were performed by J.L., L.L.H., R.B., and B.H.; the manuscript was written by J.L.; all authors commented on the final manuscript.

## References

Baker, C.S., Steel, D., Nieukirk, S. & Klinck, H. (2018) Environmental DNA (eDNA) from the wake of the whales: droplet digital PCR for detection and species identification. Frontiers in Marine Science, 5, 133.

Balasingham, K.D., Walter, R.P. & Heath, D.D. (2017) Residual eDNA detection sensitivity assessed by quantitative real-time PCR in a river ecosystem. Molecular Ecology Resources, 17, 523–532.

Barnes, M.A. & Turner, C.R. (2016) The ecology of environmental DNA and implications for conservation genetics. Conservation Genetics, 17, 1–17.

Barnes, M.A., Turner, C.R., Jerde, C.L., Renshaw, M.A., Chadderton, W.L. & Lodge, D.M. (2014) Environmental conditions influence eDNA persistence in aquatic systems. Environmental Science & Technology, 48, 1819–1827.

Bronner, I.F., Quail, M.A., Turner, D.J. & Swerdlow, H. (2014) Improved protocols for Illumina sequencing. Current Protocols in Human Genetics, 80, 18.12.11-18.12.42.

Civade, R., Dejean, T., Valentini, A., Roset, N., Raymond, J.-C., Bonin, A., … Pont, D. (2016) Spatial representativeness of environmental DNA metabarcoding signal for fish biodiversity assessment in a natural freshwater system. PloS ONE, 11, e0157366.

De Barba, M., Miquel, C., Boyer, F., Mercier, C., Rioux, D., Coissac, E. & Taberlet, P. (2014) DNA metabarcoding multiplexing and validation of data accuracy for diet assessment: application to omnivorous diet. Molecular Ecology Resources, 14, 306–323.

Deiner, K. & Altermatt, F. (2014) Transport distance of invertebrate environmental DNA in a natural river. PLoS ONE, 9, e88786.

Deiner, K., Bik, H.M., Mächler, E., Seymour, M., Lacoursière-Roussel, A., Altermatt, F., … Vere, N. (2017) Environmental DNA metabarcoding: transforming how we survey animal and plant communities. Molecular Ecology, 26, 5872–5895.

Deiner, K., Fronhofer, E.A., Machler, E., Walser, J.C. & Altermatt, F. (2016) Environmental DNA reveals that rivers are conveyer belts of biodiversity information. Nature Communications, 7, 1–9.

Dejean, T., Valentini, A., Duparc, A., Pellier-Cuit, S., Pompanon, F., Taberlet, P. & Miaud, C. (2011) Persistence of environmental DNA in freshwater ecosystems. PLoS ONE, 6, e23398.

Dunker, K.J., Sepulveda, A.J., Massengill, R.L., Olsen, J.B., Russ, O.L., Wenburg, J.K. & Antonovich, A. (2016) Potential of environmental DNA to evaluate Northern pike (*Esox lucius*) eradication efforts: an experimental test and case study. PloS ONE, 11, e0162277.

Eichmiller, J.J., Bajer, P.G. & Sorensen, P.W. (2014) The relationship between the distribution of common carp and their environmental DNA in a small lake. PloS ONE, 9, e112611.

Evans, N.T., Li, Y., Renshaw, M.A., Olds, B.P., Deiner, K., Turner, C.R., … Pfrender, M.E. (2017) Fish community assessment with eDNA metabarcoding: effects of sampling design and bioinformatic filtering. Canadian Journal of Fisheries and Aquatic Sciences, 74, 1362–1374.

Goldberg, C.S., Pilliod, D.S., Arkle, R.S. & Waits, L.P. (2011) Molecular detection of vertebrates in stream water: a demonstration using Rocky Mountain tailed frogs and Idaho giant salamanders. PLoS ONE, 6, e22746.

Goldberg, C.S., Sepulveda, A., Ray, A., Baumgardt, J. & Waits, L.P. (2013) Environmental DNA as a new method for early detection of New Zealand mudsnails (Potamopyrgus antipodarum). Freshwater Science, 32, 792–800.

Goldberg, C.S., Strickler, K.M. & Fremier, A.K. (2018) Degradation and dispersion limit environmental DNA detection of rare amphibians in wetlands: increasing efficacy of sampling designs. Science of the Total Environment, 633, 695–703.

Hänfling, B., Lawson Handley, L., Read, D.S., Hahn, C., Li, J., Nichols, P., … Winfield, I.J. (2016) Environmental DNA metabarcoding of lake fish communities reflects longterm data from established survey methods. Molecular Ecology, 25, 3101–3119.

Harper, L.R., Lawson Handley, L., Hahn, C., Boonham, N., Rees, H.C., Gough, K.C., … Hanfling, B. (2018) Needle in a haystack? A comparison of eDNA metabarcoding and targeted qPCR for detection of the great crested newt (Triturus cristatus). Ecology and Evolution, 8, 6330–6341.

Jane, S.F., Wilcox, T.M., McKelvey, K.S., Young, M.K., Schwartz, M.K., Lowe, W.H., … Whiteley, A.R. (2015) Distance, flow and PCR inhibition: eDNA dynamics in two headwater streams. Molecular Ecology Resources, 15, 216–227.

Jerde, C.L., Olds, B.P., Shogren, A.J., Andruszkiewicz, E.A., Mahon, A.R., Bolster, D. & Tank, J.L. (2016) Influence of stream bottom substrate on retention and transport of vertebrate environmental DNA. Environmental Science & Technology, 50, 8770–8779.

Kelly, R.P., Gallego, R. & Jacobs-Palmer, E. (2018) The effect of tides on nearshore environmental DNA. PeerJ, 6, e4521.

Kelly, R.P., Port, J.A., Yamahara, K.M. & Crowder, L.B. (2014) Using environmental DNA to census marine fishes in a large mesocosm. PLoS ONE, 9, e86175.

Kitson, J.J.N., Hahn, C., Sands, R.J., Straw, N.A., Evans, D.M. & Lunt, D.H. (2018) Detecting host-parasitoid interactions in an invasive Lepidopteran using nested tagging DNA-metabarcoding. Molecular Ecology, https:/doi.org/10.1111/mec.14518.

Klymus, K.E., Richter, C.A., Chapman, D.C. & Paukert, C. (2015) Quantification of eDNA shedding rates from invasive bighead carp Hypophthalmichthys nobilis and silver carp Hypophthalmichthys molitrix. Biological Conservation, 183, 77–84.

Lance, R.F., Klymus, K.E., Richter, C.A., Guan, X., Farrington, H.L., Carr, M.R., … Baerwaldt, K.L. (2017) Experimental observations on the decay of environmental DNA from bighead and silver carps. Management of Biological Invasions, 8, 343–359.

Lawson Handley, L. (2015) How will the ‘molecular revolution’ contribute to biological recording? Biological Journal of the Linnean Society, 115, 750–766.

Lawson Handley, L.J., Read, D., Winfield, I., Kimbell, H., Johnson, H., Li, J., … Haenfling, B. (2018) Temporal and spatial variation in distribution of fish environmental DNA in England’s largest lake. bioRxiv, https://doi.org/10.1101/376400.

Li, J., Lawson Handley, L.J., Read, D.S. & Hänfling, B. (2018a) The effect of filtration method on the efficiency of environmental DNA capture and quantification via metabarcoding. Molecular Ecology Resources, 18, 1102–1114.

Li, Y., Evans, N.T., Renshaw, M.A., Jerde, C.L., Olds, B.P., Shogren, A.J., … Pfrender, M.E. (2018b) Estimating fish alpha- and beta-diversity along a small stream with environmental DNA metabarcoding. Metabarcoding and Metagenomics, 2, e24262.

Maruyama, A., Nakamura, K., Yamanaka, H., Kondoh, M. & Minamoto, T. (2014) The release rate of environmental DNA from juvenile and adult fish. PLoS ONE, 9, e114639.

O’Donnell, J.L., Kelly, R.P., Shelton, A.O., Samhouri, J.F., Lowell, N.C. & Williams, G.D. (2017) Spatial distribution of environmental DNA in a nearshore marine habitat. PeerJ, 5, e3044.

Oksanen, J., Blanchet, F.G., Friendly, M., Kindt, R., Legendre, P., McGlinn, D., … Wagner, H. (2017) Vegan: community ecology package. Retrieved from https://CRAN.R-project.org/package=vegan.

Pilliod, D.S., Goldberg, C.S., Arkle, R.S. & Waits, L.P. (2013) Estimating occupancy and abundance of stream amphibians using environmental DNA from filtered water samples. Canadian Journal of Fisheries and Aquatic Sciences, 70, 1123–1130.

Pilliod, D.S., Goldberg, C.S., Arkle, R.S. & Waits, L.P. (2014) Factors influencing detection of eDNA from a stream-dwelling amphibian. Molecular Ecology Resources, 14, 109–116.

Port, J.A., O’Donnell, J.L., Romero-Maraccini, O.C., Leary, P.R., Litvin, S.Y., Nickols, K.J., … Kelly, R.P. (2016) Assessing vertebrate biodiversity in a kelp forest ecosystem using environmental DNA. Molecular Ecology, 25, 527–541.

R_Core_Team (2018) R: a language and environment for statistical computing. R Foundation for Statistical Computing, Vienna, Austria. Retrieved from http://www.R-project.org.

Rees, H.C., Maddison, B.C., Middleditch, D.J., Patmore, J.R. & Gough, K.C. (2014) The detection of aquatic animal species using environmental DNA–a review of eDNA as a survey tool in ecology. Journal of Applied Ecology, 51, 1450–1459.

Riaz, T., Shehzad, W., Viari, A., Pompanon, F., Taberlet, P. & Coissac, E. (2011) ecoPrimers: inference of new DNA barcode markers from whole genome sequence analysis. Nucleic Acids Research, 39, e145.

Sassoubre, L.M., Yamahara, K.M., Gardner, L.D., Block, B.A. & Boehm, A.B. (2016) Quantification of environmental DNA (eDNA) shedding and decay rates for three marine fish. Environmental Science & Technology, 50, 10456–10464.

Sato, H., Sogo, Y., Doi, H. & Yamanaka, H. (2017) Usefulness and limitations of sample pooling for environmental DNA metabarcoding of freshwater fish communities. Scientific Reports, 7, 14860.

Seymour, M., Durance, I., Cosby, B.J., Ransom-Jones, E., Deiner, K., Ormerod, S.J., … de Bruyn, M. (2018) Acidity promotes degradation of multi-species environmental DNA in lotic mesocosms. Communications Biology, 1, 4.

Shogren, A.J., Tank, J.L., Andruszkiewicz, E., Olds, B., Mahon, A.R., Jerde, C.L. & Bolster, D. (2017) Controls on eDNA movement in streams: Transport, Retention, and Resuspension. Scientific Reports, 7, 5065.

Shogren, A.J., Tank, J.L., Andruszkiewicz, E.A., Olds, B., Jerde, C. & Bolster, D. (2016) Modelling the transport of environmental DNA through a porous substrate using continuous flow-through column experiments. Journal of the Royal Society Interface, 13, 20160290.

Spear, S.F., Groves, J.D., Williams, L.A. & Waits, L.P. (2015) Using environmental DNA methods to improve detectability in a hellbender (*Cryptobranchus alleganiensis*) monitoring program. Biological Conservation, 183, 38–45.

Stéphane, D., Anne-Béatrice, D. & Jean, T. (2018) ade4: analysis of ecological data: exploratory and euclidean methods in environmental sciences. Retrieved from https://CRAN.R-project.org/package=ade4. Retrieved from https://CRAN.R-project.org/package=ade4.

Stoeckle, B.C., Beggel, S., Cerwenka, A.F., Motivans, E., Kuehn, R. & Geist, J. (2017) A systematic approach to evaluate the influence of environmental conditions on eDNA detection success in aquatic ecosystems. PloS ONE, 12, e0189119.

Strickler, K.M., Fremier, A.K. & Goldberg, C.S. (2015) Quantifying effects of UV-B, temperature, and pH on eDNA degradation in aquatic microcosms. Biological Conservation, 183, 85–92.

Taberlet, P., Coissac, E., Hajibabaei, M. & Rieseberg, L.H. (2012) Environmental DNA. Molecular Ecology, 21, 1789–1793.

Takahara, T., Minamoto, T., Yamanaka, H., Doi, H. & Kawabata, Z.i. (2012) Estimation of fish biomass using environmental DNA. PLoS ONE, 7, e35868.

Thomsen, P.F., Kielgast, J., Iversen, L.L., Møller, P.R., Rasmussen, M. & Willerslev, E. (2012a) Detection of a diverse marine fish fauna using environmental DNA from seawater samples. PLoS ONE, 7, e41732.

Thomsen, P.F., Kielgast, J., Iversen, L.L., Wiuf, C., Rasmussen, M., Gilbert, M.T.P., … Willerslev, E. (2012b) Monitoring endangered freshwater biodiversity using environmental DNA. Molecular Ecology, 21, 2565–2573.

Thomsen, P.F. & Willerslev, E. (2015) Environmental DNA–an emerging tool in conservation for monitoring past and present biodiversity. Biological Conservation, 183, 4–18.

Tillotson, M.D., Kelly, R.P., Duda, J.J., Hoy, M., Kralj, J. & Quinn, T.P. (2018) Concentrations of environmental DNA (eDNA) reflect spawning salmon abundance at fine spatial and temporal scales. Biological Conservation, 220, 1–11.

Turner, C.R., Uy, K.L. & Everhart, R.C. (2015) Fish environmental DNA is more concentrated in aquatic sediments than surface water. Biological Conservation, 183, 93–102.

Valentini, A., Taberlet, P., Miaud, C., Civade, R., Herder, J., Thomsen, P.F., … Dejean, T. (2016) Next-generation monitoring of aquatic biodiversity using environmental DNA metabarcoding. Molecular Ecology, 25, 929–942.

Wickham, H. & Chang, W. (2016) ggplot2: create elegant data visualisations using the grammar of graphics. Retrieved from https://CRAN.R-project.org/package=ggplot2.

Wilcox, T.M., McKelvey, K.S., Young, M.K., Sepulveda, A.J., Shepard, B.B., Jane, S.F., … Schwartz, M.K. (2016) Understanding environmental DNA detection probabilities: a case study using a stream-dwelling char Salvelinus fontinalis. Biological Conservation, 194, 209–216.

